## Supplementary material for "TNFα-induced LDL cholesterol accumulation involve elevated LDLR cell surface levels and SR-B1 downregulation in human arterial endothelial cells": Original blots

### Original blots to Figure 6

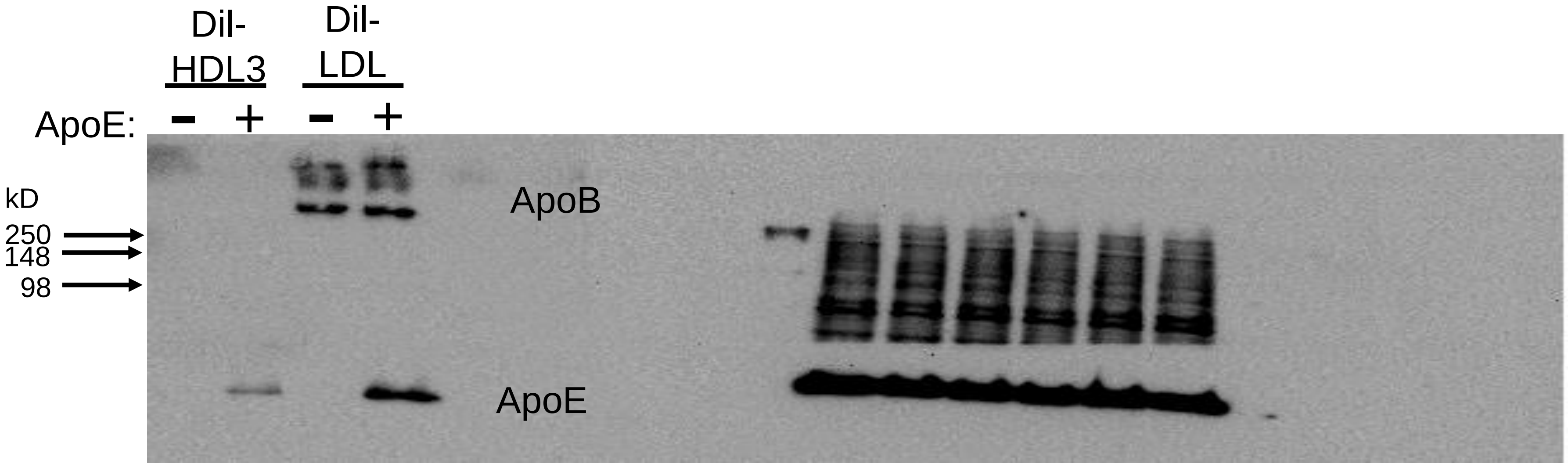


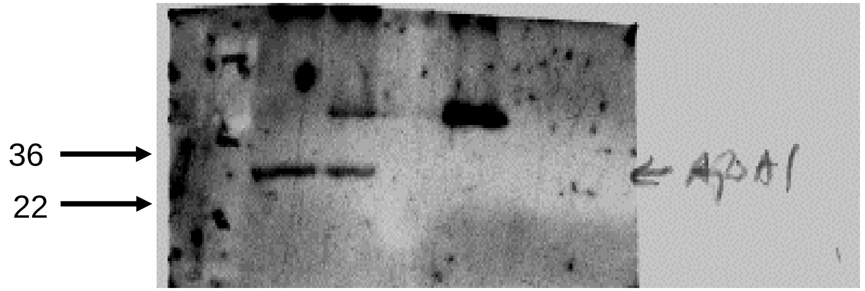


ApoA1,
Post ApoB and apoE

### Original blots to Figure 7 from the same lysates

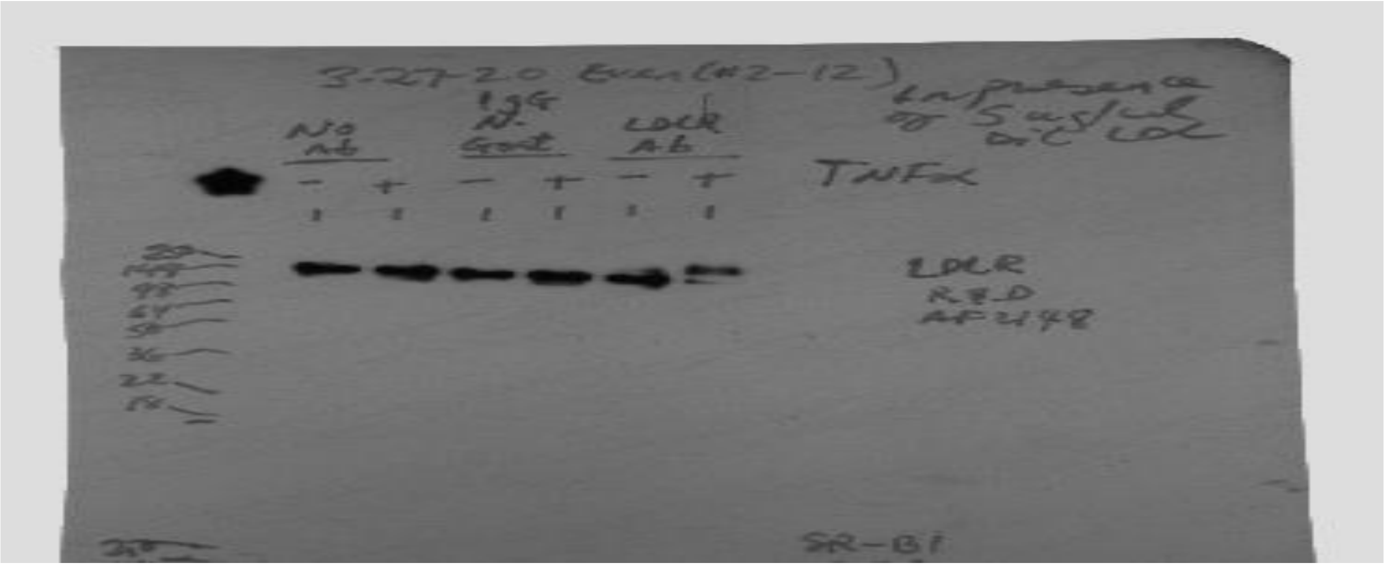


LDLR

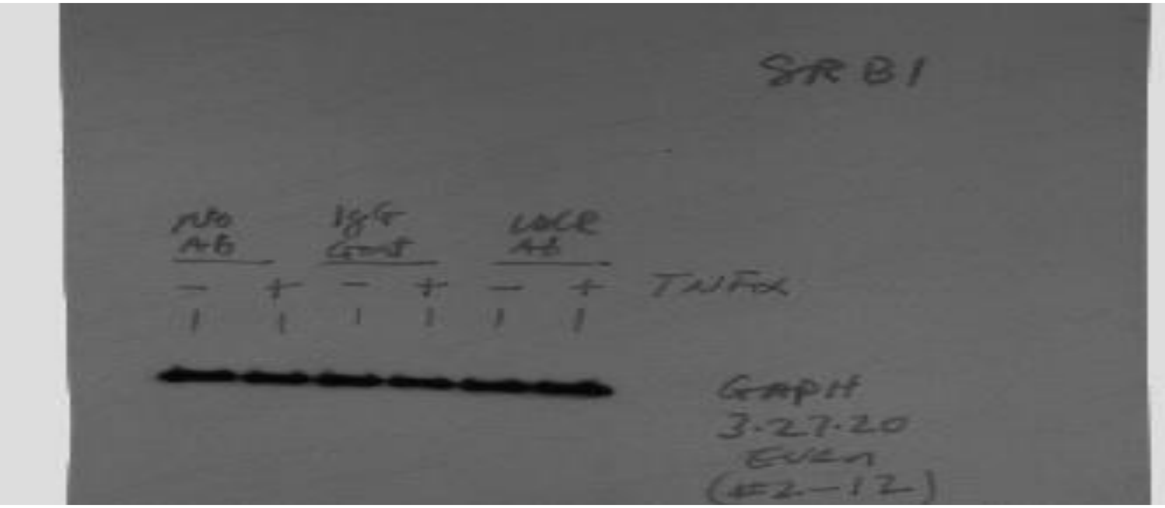


GAPDH

**Original blots to Figure 8 from the same lysates**

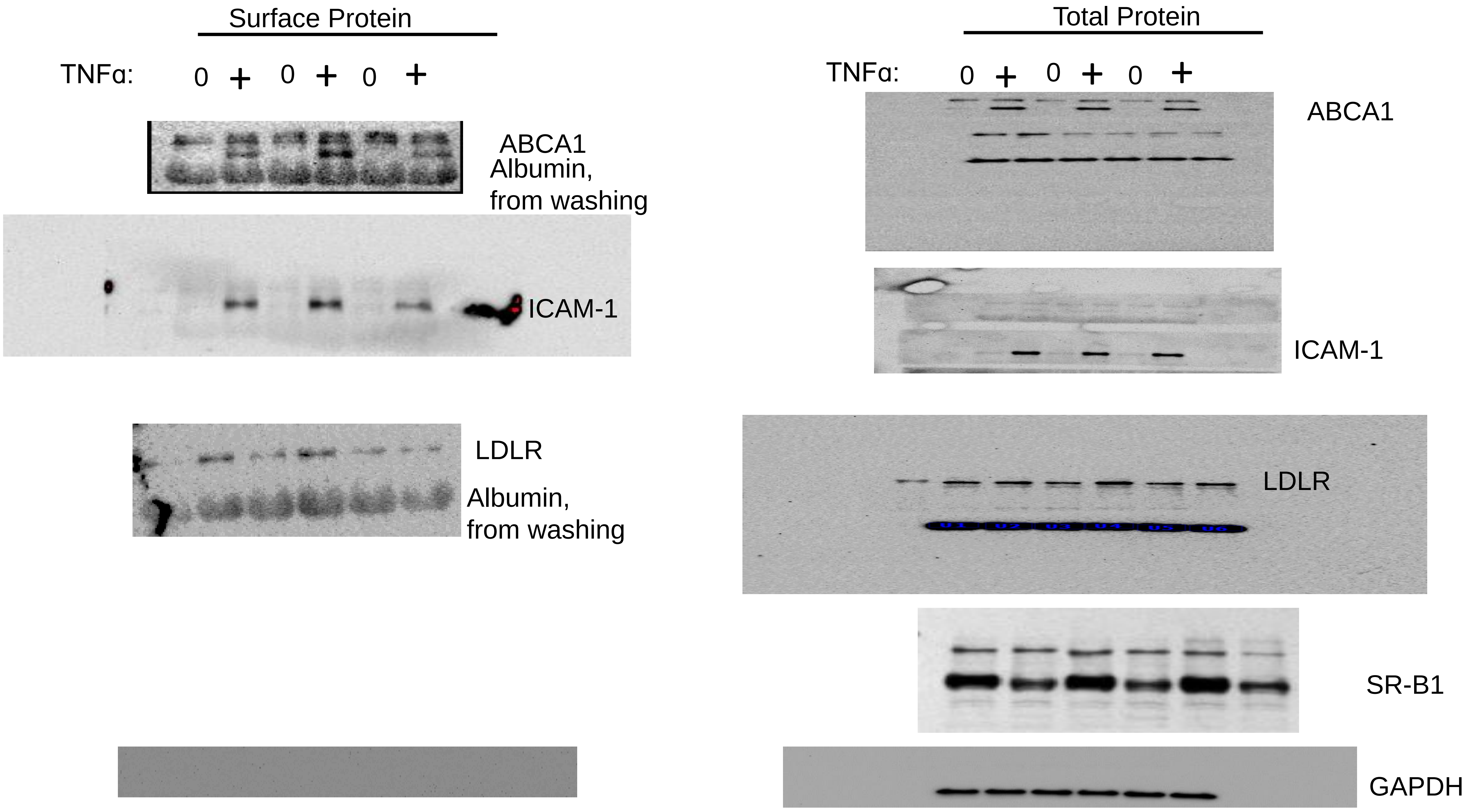
